# Acquisition of specific human respiratory tract binding by 2.3.4.4b H5N1 hemagglutinins requires multiple mutations

**DOI:** 10.64898/2026.04.16.718875

**Authors:** María Ríos Carrasco, Mafalda F. Guerreiro Cabana, Eszter Kovács, Zoé Ducarne, Geert-Jan Boons, Robert P. de Vries

## Abstract

It has been suggested that the hemagglutinin of the human-infecting cattle-derived 2.3.4.4b virus A/Texas/34 (H5TX) requires only one mutation, namely Q226L, to switch from binding avian-type to human-type receptor preference. In this study, we examined the binding of H5TX Q226L, along with other key mutations, to sections of human trachea. We conclude that, while H5TX Q226L can bind human-type receptors, more than a single mutation is required for this protein to bind to human respiratory tract tissue. We also report changes in receptor-binding specificity of another 2.3.4.4b HA mutant, H5FR Q226L, associated with the presence of a multibasic cleavage site. This study offers insight into the determinants of evolution towards human-type receptor binding in currently circulating H5Nx viruses. It also emphasizes the importance of testing individual strains using additional methods, including tissue-based approaches, alongside synthetic glycans.

**Importance:** Currently, H5N1 influenza A viruses are responsible for numerous zoonotic spillover events, from infecting birds to other mammals, including dairy cattle. Although no human-to-human transmission has been observed, several people have been infected. This host range expansion is typically linked to changes in one of the viral surface proteins, hemagglutinin, which can switch its preference from avian-type to human-type receptors. To better understand the potential of the currently circulating H5N1 virus to transmit among humans, we evaluated the effects of the Q226L mutation, in combination with other amino acid substitutions, on binding to the human trachea. We also studied the effect of the multibasic cleavage site, a specific motif present in highly pathogenic influenza strains, on receptor binding properties. These findings provide insight into the role of receptor binding in influenza infections.

## Introduction

Highly pathogenic avian influenza (HPAI) H5 viruses of the 2.3.4.4 lineage continue to pose a growing zoonotic threat as they expand into new host species, leading to increasingly frequent spillover events (1, 2). These viruses are typically adapted to transmit among avian species because they prefer avian-type receptors, namely α2,3-linked sialic acids (SIA), which are abundant in birds (3-6). However, recent outbreaks in dairy cattle, together with sporadic human infections, have raised concerns about the acquisition of a broader receptor-binding profile than that of classical avian influenza A viruses (6-8).

Literature highlights that only minimal genetic changes in the hemagglutinin (HA) of these circulating viruses may be required to switch binding from avian-type receptors, α2,3-linked SIA, to human-type receptors, α2,6-linked SIA (9). Two recent studies demonstrate that a single amino acid substitution, Q226L, in distinct H5 HA backgrounds can shift glycan binding towards α2,6-linked SIA (10, 11). Structural and glycan-binding analyses reveal that Q226L remodels the receptor-binding site (RBS) to promote human-type receptor engagement, although additional HA features modulate the ease and extent of this shift depending on the antigenic background (12-15). However, more extended analyses of binding to human upper respiratory tissues are still lacking.

Although receptor specificity is usually governed by residues in the RBS and adjacent loops, the effect of the presence or absence of a multibasic/polybasic cleavage site (MBCS) on HA receptor-binding properties remains unknown. HAs with a MBCS are cleaved by ubiquitous furin-like host proteases, allowing systemic infection in poultry. Previously, experimental insertions of an MBCS into human HAs have been reported to typically interfere with replication or affect virulence, but do not, on their own, change receptor specificity (16, 17). However, changes in HA processing, trafficking, protein stability, and fusion conditions due to the acquisition of an MBCS can indirectly influence which cell types are infected and, thus, the apparent receptor usage in vivo (17). For instance, some studies show that combinations of MBCS and other HA mutations (including those in HA2 or adjacent domains) can indeed affect H7 tropism or virulence (18, 19).

In this study, we assess the receptor-binding profile of a representative virus currently circulating among cows, A/Texas/37/24 H5 HA (H5TX), based on key mutations in and around the RBS. Our observations revealed that although H5TX Q226L shows a shift towards preferring α2,6-linked sialosides, the switch is observed only for elongated and complex glycans and does not engage receptors present in the human trachea. These findings align with unsustained human-to-human transmission, given the lack of attachment to the upper respiratory tract as a first stage of infection.

We also addressed the MBCS knowledge gap in this paper by inserting or removing the MBCS sequence in several HAs from different antigenic backgrounds, including multiple H5 strains with their respective Q226L mutants, as well as a human H1N1 HA. Hemagglutination, tissue staining, glycan array, and glycan-based ELISA results indicated that the presence of MBCS did not affect receptor-binding specificity, except for H5FR Q226L. The insertion of an MBCS in H5FR Q226L resulted in the loss of the characteristic dual receptor binding previously reported for this mutant (10).

These findings demonstrate the importance of assessing each antigenic strain individually to understand how single mutations and HA changes affect its structural context and alter receptor specificity.

## Results

### H5TX Q226L fails to bind the human trachea

It has been reported that A/Texas/37/24 H5 HA (H5TX WT) only requires a single mutation, Q226L, to switch its receptor-binding specificity from avian to human-type receptors (11). In this study, we expressed H5TX and its Q226L mutant (H5TX Q226L) HAs, and tested their binding profiles using a previously published glycan array (20, 21), which comprises a library of compounds, including non-sialylated, α2,3-, and α2,6-linked sialic acids with one to four LacNac repeats (Figure 1A). H5TX WT bound to α2,3-linked compounds, as expected for an avian H5 HA, and previously described (22, 23). In the case of H5TX Q226L, binding to α2,3-linked sialosides was completely lost and switched to very weak binding to α2,6-linked tri- and one tetra-LacNac compound **(#18**) (Figure 1A).

**Figure 1:**
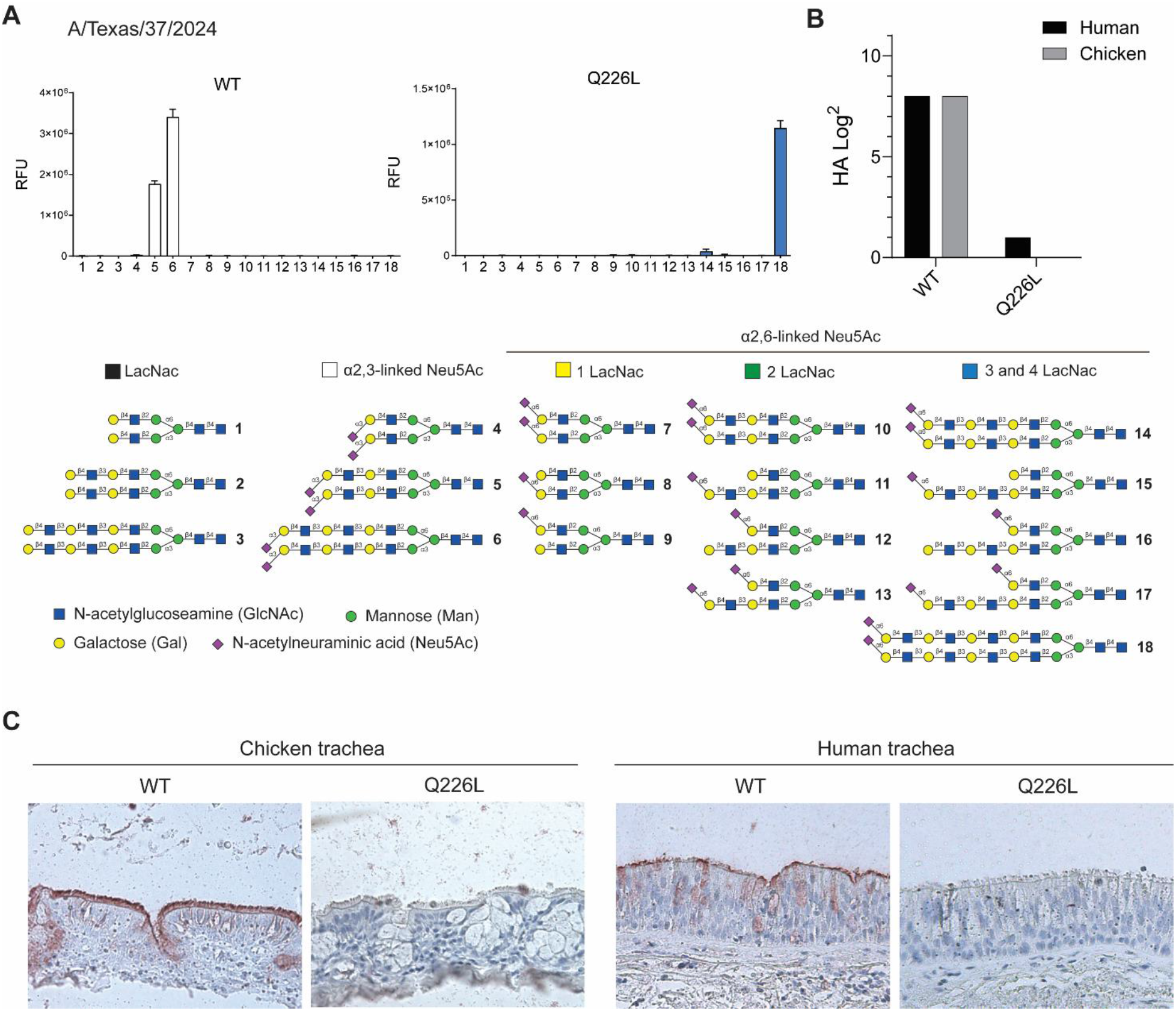
(A) The glycan microarray was used to determine the receptor specificities of A/Texas/37/2024 WT and Q226L HAs. An overview of the synthetic glycans printed on the microarray (n = 6) is shown. For each glycan, the mean signal and standard deviation were calculated from six replicates. The data are representative of three independent assays performed using antibody-precomplexed protein solutions. Glycans 1–3 are non-sialylated controls (black bars), α2,3-linked N-acetyl glycans are shown in white (glycans 4–6), and α2,6-linked N-acetyl glycans are displayed as yellow bars for one LacNAc repeat (glycans 7–9), green bars for two LacNAc repeats (glycans 10–13), and blue bars for three and four LacNAc repeats (glycans 14–18). (B) Hemagglutination assay using human and chicken erythrocytes with A/Texas/37/2024 WT and Q226L mutant HAs. (C) The binding specificity of A/Texas/37/2024 WT and Q226L HAs was evaluated by immunohistochemical staining of chicken and human trachea. Tissue sections were visualized using AEC staining.

To broaden our receptor display, we examined the binding of these HAs to chicken erythrocytes, which are abundant in α2,3-linked SIA, and human erythrocytes, which mostly display α2,6-linked SIA. H5TX WT was able to bind both, while only very weak binding was observed to human erythrocytes for H5TX Q226L (Figure 1B).

The binding of HA to upper respiratory tract tissue sections has been extensively linked to the virus’s ability to infect certain hosts. We used chicken and human tracheal sections to assess the binding of our H5TX HAs to avian- and human-type receptors in the upper respiratory tract. Interestingly, we observe binding of H5TX WT to chicken and human trachea, albeit to a lesser extent in the latter, matching the broadened receptor-binding profile for α2,3-linked SIA on mucin-like O-glycans of 2.3.4.4b H5 proteins we previously described (24) (Figure 1C). When we analyze the H5TX Q226L, binding is abrogated in both chicken and human trachea (Figure 1C).

In summary, glycan array studies indicate that H5TX Q226L switches binding from α2,3-linked to elongated α2,6-linked SIAs. However, tissue immunohistochemical staining confirms that these complex glycans, containing three or four LacNac repeats, are absent from the human upper respiratory tract, as previously reported (25). This observation calls into question whether Q226L is the only HA mutation required for H5TX to fully adapt to human-type receptors, as previously suggested.

### H5TX N224K Q226L rescues binding to human trachea sections

Previous research reported that H5TX Q226L was unable to engage with linear α2,6-linked SIA. When combined with G228S, a mutation that often co-evolves with Q226L, binding to these compounds was not observed. However, H5TX N224K Q226L, another previously reported combination (13), bound to 6-SLN_3_. We inserted N224K into our H5TX Q226L HA and tested its binding to tissue sections. While it did not bind chicken trachea, H5TX Q226L N224K recovered binding to human trachea, similarly to H5TX WT (Figure 2A). On our glycan array, this protein showed a preference for α2,6-linked biantennary glycans with three and four LacNac repeats, some of which are asymmetrically sialylated **(#17**) (Figure 2B) (20, 21).

**Figure 2:**
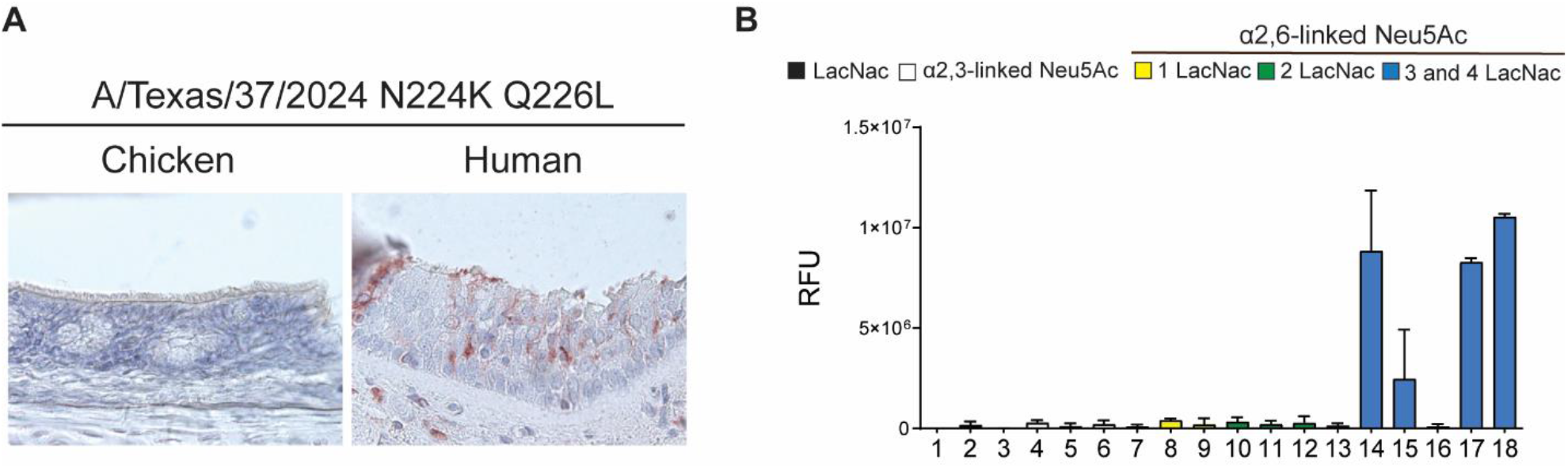
(A) The binding specificity of A/Texas/37/2024 Q226L N224K HA was evaluated by immunohistochemical staining of chicken and human trachea. Tissue sections were visualized using AEC staining. (B) The glycan microarray was used to determine the receptor specificity of A/Texas/37/2024 Q226L N224K HA. See Figure 1 for an overview of the synthetic glycans printed on the microarray (n = 6). For each glycan, the mean signal and standard deviation were calculated from six replicates. The data are representative of three independent assays performed using antibody-precomplexed protein solutions. Glycans 1–3 are non-sialylated controls (black bars), α2,3-linked N-acetyl glycans are shown in white (glycans 4–6), and α2,6-linked N-acetyl glycans are displayed as yellow bars for one LacNAc repeat (glycans 7–9), green bars for two LacNAc repeats (glycans 10–13), and blue bars for three and four LacNAc repeats (glycans 14– 18).

### The Q226L mutation is not sufficient for the H5TX HA to fully switch binding to α2,6-linked SIA

The HA protein derived from A/black swan/Akita/1/16 (H5N6), belonging to the 2.3.4.4e clade (H5AK), is able to fully switch receptor binding preference from avian to human-type receptors by introducing the Q226L mutation (10). However, while H5AK Q226L binds avidly to human trachea, H5TX Q226L does not (Figure 1C), suggesting that additional mutations are needed to fully adapt to receptors present in the human upper respiratory tract. Two residues in the RBS abrogated α2,6-linked SIA binding in the H5AK Q226L background, namely a leucine deletion in position 133 (Δ133L) and glutamine in position 227 (Q227). We introduced these residues individually and in combination with the Q226L mutation in the H5TX background. We also made the triple mutant H5TX Δ133L Q226L R227Q to resemble the H5AK Q226L sequence.

H5TX Δ133L and R227Q single mutants bound to both chicken and human trachea, suggesting that none of these make an impact on H5TX receptor binding individually (Figure 3A). Nevertheless, when these mutations are introduced together with Q226L, binding is completely lost, as observed for the single H5TX Q226L mutant. This loss of binding to both human and chicken trachea was also observed for the double mutants (H5TX Δ113L Q226L and Q226L R227Q), as well as for the triple mutant (H5TX Δ133L Q226L R227Q) (Figure 3A).

**Figure 3:**
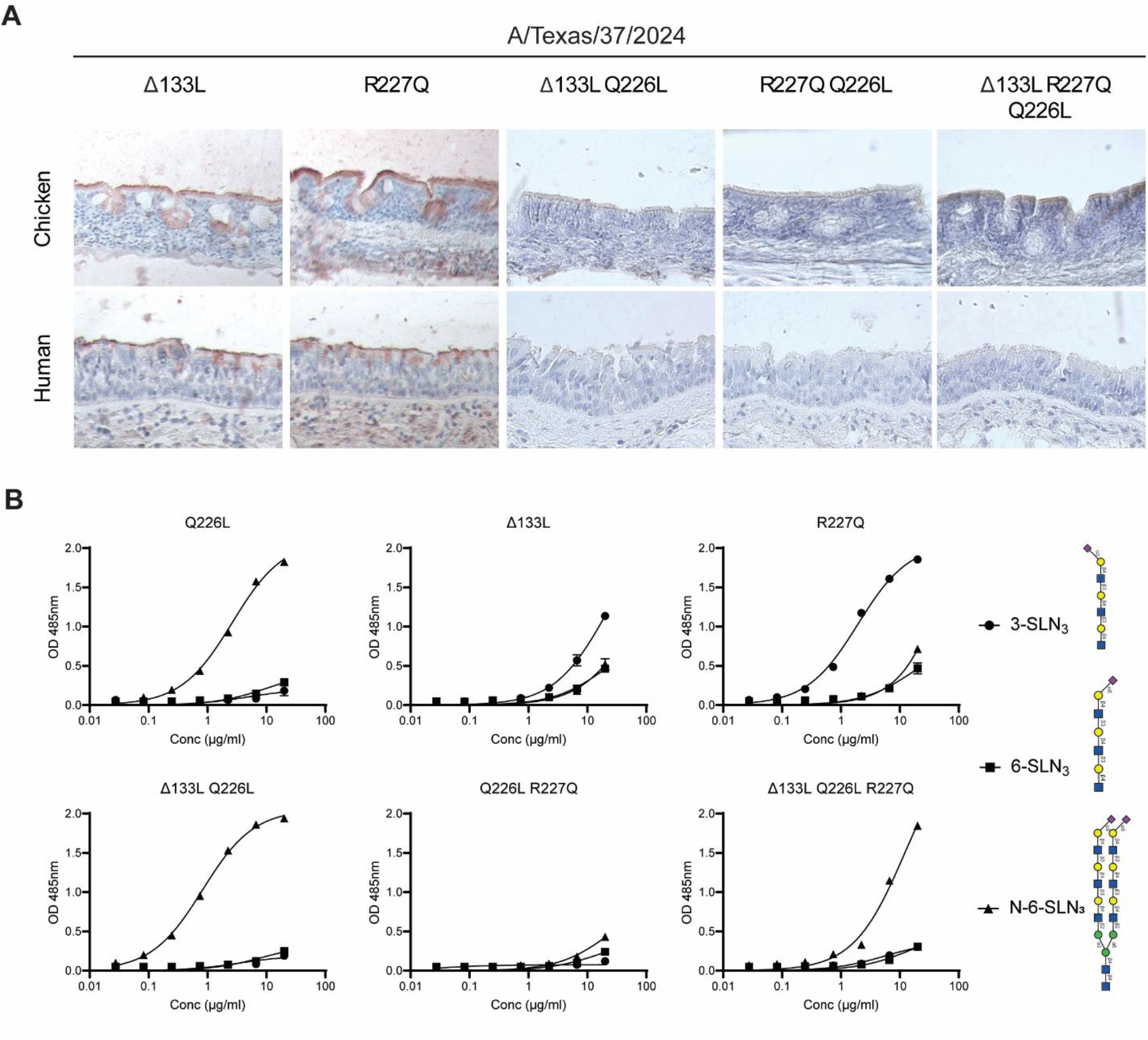
(A) The binding specificity of A/Texas/37/2024 Δ133L, R227Q, Δ133L Q226L, R227Q Q226L, and Δ133L R227Q Q226L mutant HAs was evaluated by immunohistochemical staining of chicken and human trachea. Tissue sections were visualized using AEC staining. (B) The binding specificities of A/Texas/37/2024 Δ133L, R227Q, Δ133L Q226L, R227Q Q226L, and Δ133L R227Q Q226L mutant HAs were further evaluated by glycan ELISA. Biotinylated linear triLacNac α2,3-linked SIA (3-SLN_3_), linear triLacNac α2,6-linked SIA (6-SLN_3_), and biantennary triLacNac α2,6-linked SIA (N-6-SLN_3_) were coated on streptavidin-coated plates. Antibody precomplexed protein mixtures were added, and binding was detected by adding HRP substrate and measured by absorbance at 485nm.

We tested these mutant HA proteins in a glycan ELISA-based assay, using linear triLacNac with both α2,3-linked SIA (3-SLN_3_) and α2,6-linked SIA (6-SLN_3_), and a biantennary triLacNac α2,6-linked SIA compound (N-6-SLN_3_) (Figure 3B). H5TX WT binding to 3-SLN_3_ was conserved in H5TX Δ133L and H5TX R227Q, consistent with our observation of tissue binding in chicken trachea. Neither H5TX WT nor any of its mutants were able to engage with 6-SLN_3_. H5TX Q226L bound to N-6-SLN_3_, as previously reported (11). We did not observe binding to N-6-SLN_3_ by the single mutants H5TX Δ133L and H5TX R227Q. H5TX Δ133L Q226L binding to N-6-SLN_3_ suggests that Q226L is the driver of the preference for this compound. This N-6-SLN_3_ binding is completely abrogated by R227Q in H5TX Q226L R227Q, but partially recovered in the triple mutant H5TX Δ133L Q226L.

In summary, the combination of Δ133L and R227Q in H5TX Q226L did not result in the binding phenotype observed for H5AK Q226L, where there is a full switch from 3-SLN_3_ to 6-SLN_3_, supported by binding to human trachea sections (10). In the case of H5TX mutants, the receptor-binding switch is observed only for N-6-SLN_3_, which, based on the lack of binding to human trachea, is not present in the human upper respiratory tract.

### The multibasic cleavage site influences receptor binding in a background-dependent manner

H5FR (from H5/duck/France/161108/16) is the evolutionary precursor of H5TX and has shown dual binding to chicken and human trachea (10). Furthermore, when the Q226L mutation is introduced, it conserves binding to α2,3-linked sialosides (10). H5FR and H5TX HAs only differ in six amino acid residues (L111M, L122Q, T144A, T199I, V214A, and N240D). Another difference is that the H5FR HA had its multibasic cleavage site (MBCS) sequence removed, whereas H5TX in the current study contains an MBCS. Since the effects of MBCS on these HAs’ receptor-binding properties have not been reported, we generated H5FR and H5TX, and their corresponding Q226L mutants, in the presence and absence of MBCS to study their glycan-binding profiles.

H5FR (-)MBCS and (+)MBCS kept a similar binding profile to α2,3-linked sialosides on the glycan array (Figure 4A). In contrast, while H5FR Q226L (-)MBCS maintained binding to α2,3-linked SIA and gained binding to complex α2,6-linked compounds, H5FR Q226L (+)MBCS lost α2,3-linked SIA binding completely, and switched its preference to bind elongated α2,6-linked compounds (Figure 4A). In the case of H5TX, both WT and Q226L variants maintained the same specificity regardless of the presence of the MBCS sequence.

**Figure 4:**
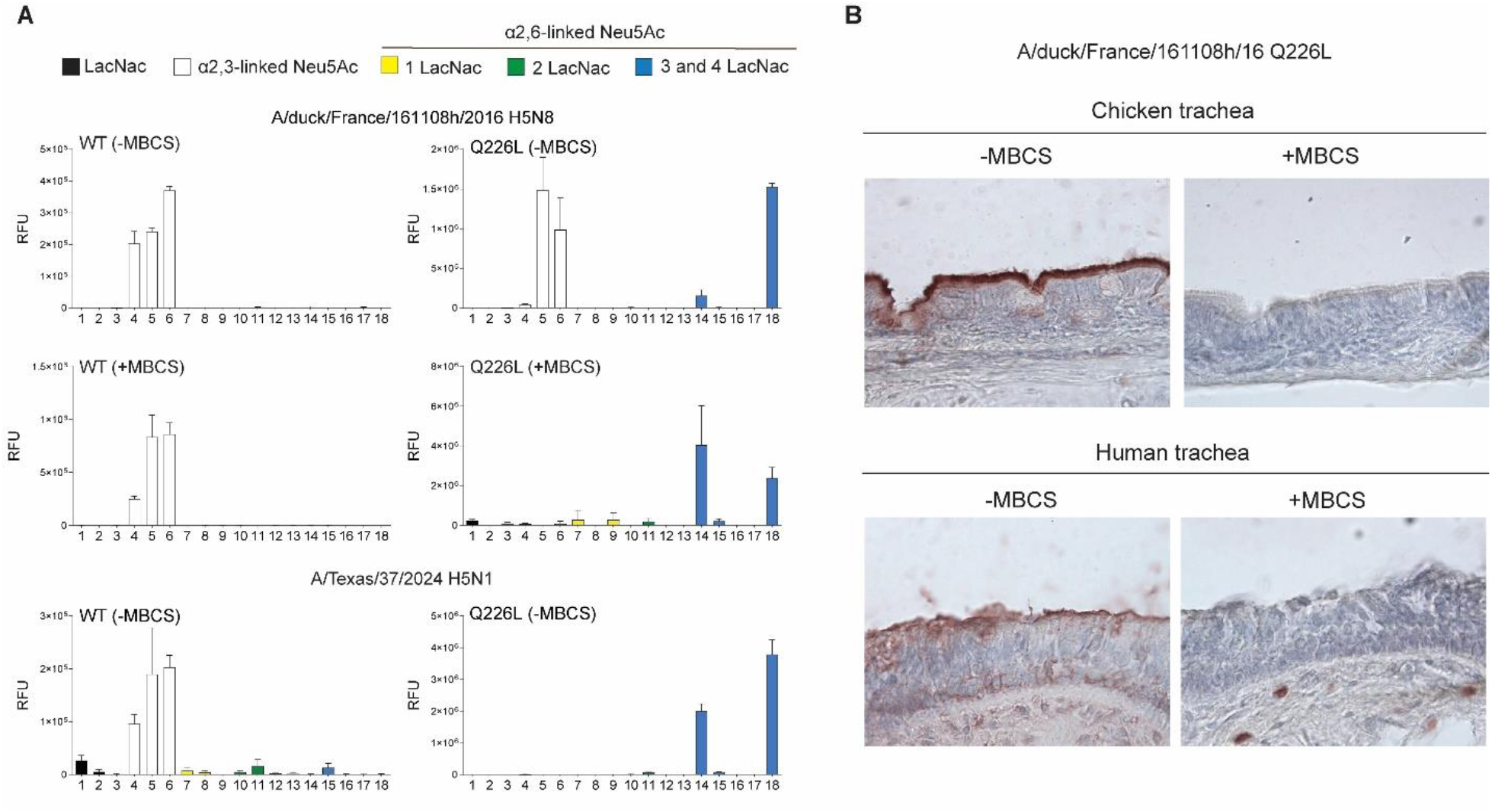
(A) The glycan microarray was used to determine the receptor specificity of A/duck/France/161108/16 WT and Q226L (+/-MBCS) and A/Texas/37/2024 WT and Q226L (+MBCS) HAs. See Figure 1 for an overview of the synthetic glycans printed on the microarray (n = 6). For each glycan, the mean signal and standard deviation were calculated from six replicates. The data are representative of three independent assays performed using antibody-precomplexed protein solutions. Glycans 1–3 are non-sialylated controls (black bars), α2,3-linked N-acetyl glycans are shown in white (glycans 4–6), and α2,6-linked N-acetyl glycans are displayed as yellow bars for one LacNAc repeat (glycans 7–9), green bars for two LacNAc repeats (glycans 10–13), and blue bars for three and four LacNAc repeats (glycans 14–18). (B) The binding specificity of A/duck/France/161108/16 Q226L (+/-MBCS) was evaluated by immunohistochemical staining of chicken and human trachea. Tissue sections were visualized using AEC staining.

In tissue immunostaining, H5FR WT (+/-)MBCS bound comparably to chicken and human trachea sections, illustrating the previously reported broad binding spectrum of this protein (24). When introducing Q226L, binding to both chicken and human trachea was lost completely in the (+)MBCS version (Figure 4B). These observations align with the glycan array data. H5TX WT(+/-)MBCS proteins were able to engage receptors in chicken and, to a lesser extent, in human trachea. The Q226L mutation abrogated protein binding in both (+) and (-) MBCS versions.

We further tested these proteins in a glycan ELISA-based assay, again with 3-SLN_3_, 6-SLN_3_, and N-6-SLN_3_ (Figure 5). H5FR WT (-/+) MBCS bound similarly to 3-SLN_3_. H5FR Q226L (-) MBCS maintained this binding to 3-SLN_3_ but could not engage with 6-SLN_3_ receptors, as previously reported (10). Although H5FR Q226L (+)MBCS could not bind any of the linear compounds, both H5FR Q226L (+/-) MBCS proteins displayed binding to N-6-SLN_3_, matching our observations on the glycan array. H5TX (+)MBCS bound slightly stronger than its (-) MBCS counterpart to 3-SLN_3_, while no binding was observed to α2,6-linked compounds. Nevertheless, H5TX Q226L (+)MBCS showed higher avidity for N-6-SLN_3_ than its (-)MBCS counterpart.

**Figure 5:**
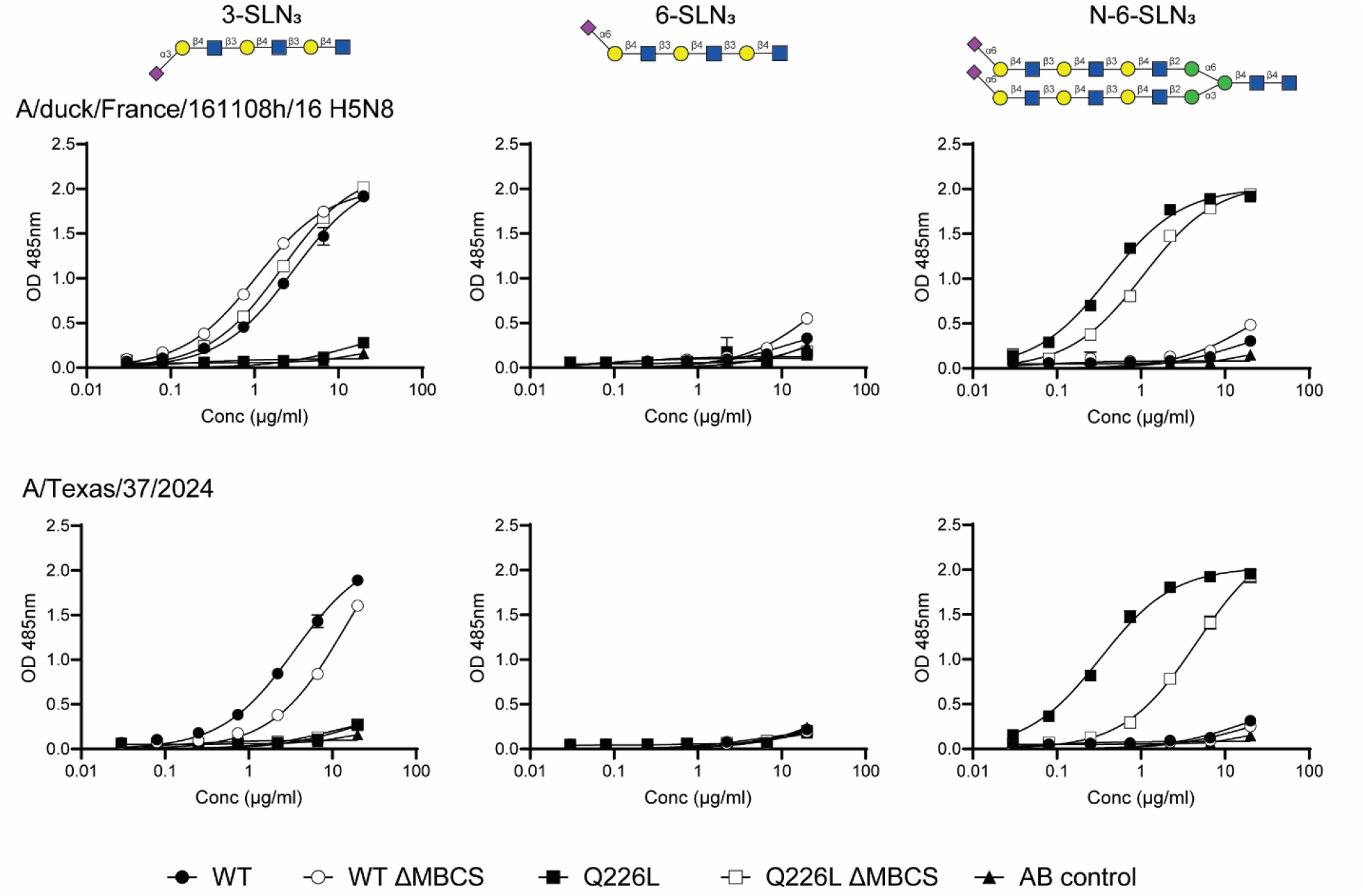
The binding specificities of A/duck/France/161108/16 WT and Q226L (+/-MBCS) and A/Texas/37/2024 WT and Q226L (+/-MBCS) HAs were further evaluated by glycan ELISA. Biotinylated linear triLacNac α2,3-linked SIA (3-SLN_3_), linear triLacNac α2,6-linked SIA (6-SLN_3_), and biantennary triLacNac α2,6-linked SIA (N-6-SLN_3_) were coated on streptavidin-coated plates. Antibody precomplexed protein mixtures were added, and binding was detected by adding HRP substrate and measured by absorbance at 485nm.

**Figure 5:**
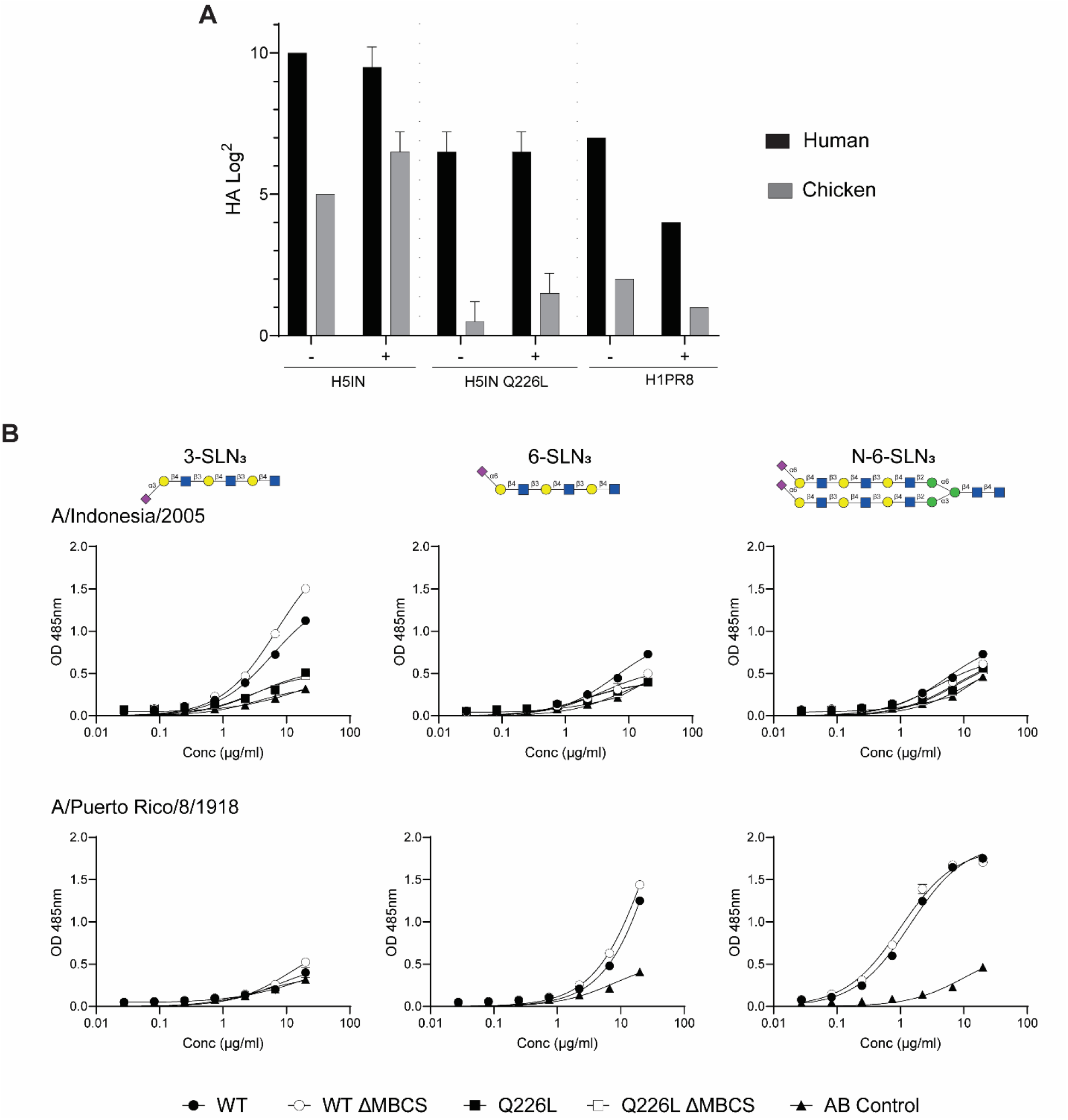
(A) Hemagglutination assay using human and chicken erythrocytes with A/Indonesia/2005 WT and Q226L (+/-MBCS) and A/Puerto Rico/8/1934 WT (+/-MBCS) HAs. (B) The binding specificities of A/Indonesia/2005 WT and Q226L (+/-MBCS) and A/Puerto Rico/8/1934 WT (+/-MBCS) HAs were further evaluated by glycan ELISA. Biotinylated linear triLacNac α2,3-linked SIA (3-SLN_3_), linear triLacNac α2,6-linked SIA (6-SLN_3_), and biantennary triLacNac α2,6-linked SIA (N-6-SLN_3_) were coated on streptavidin-coated plates. Antibody precomplexed protein mixtures were added, and binding was detected by adding HRP substrate and measured by absorbance at 485nm.

### The insertion of a multibasic cleavage site across different strains does not affect HA receptor-binding properties

To expand the study of the MBCS effect on receptor binding properties, we introduced the MBCS sequence into a low-pathogenic H5 strain from an older clade, A/Indonesia/05/2005 H5N1 (H5IN), and generated its corresponding Q226L variant, with and without MBCS. We also inserted an MBCS sequence in a human isolate HA from A/Puerto Rico/8/1934 H1N1 (H1PR8).

No changes in the hemagglutination activity of either H5IN, H5IN Q226L, and H1PR8 proteins were observed (Figure 6A). The erythrocyte-binding profile of these proteins was maintained in human and chicken blood, although a loss of avidity was noticeable upon introducing the MBCS in H1PR8.

In contrast to our observation for H5FR Q226L, receptor specificity was unaffected regardless of the presence of MBCS. Glycan ELISA results displayed similar binding profiles for H5IN WT, H5IN Q226L, and H1PR8 (Figure 6B). This suggests that the change in receptor-binding specificity of H5FR Q226L is specific to that background and mutant.

## Discussion

In this study, we investigate the shift of a representative human-derived H5N1 strain, H5TX, towards binding human-type receptors. Our results confirm this shift, although we only observe it for elongated complex N-linked glycans. We aimed to correlate these results with human trachea binding, the native site of infection. H5TX Q226L could not engage receptors in the human trachea, illustrating that the complex glycans used in the array and ELISA are not displayed in the human upper respiratory tract. Based on the literature, we explored key mutations in combination with Q226L that could retrieve human trachea binding.

Similar to previous studies with 2.3.4.4b H5FR, H5TX WT bound human tracheal tissues, despite not binding human-type receptors on the glycan array (10). This is consistent with previous observations that these proteins have a more promiscuous receptor-binding profile (24, 26). The binding of these HAs is not restricted to N-glycans; they also bind O-glycans (24). These glycans are present in the human upper respiratory tract; hence, avian strains bind to the human trachea (27).

Based on the literature, H5TX N224K Q226L bound simple linear α2,6-linked SIA in ELISA and surface plasmon resonance (11). This N224K mutation has already been reported to contribute to the transmissibility among ferrets for A/Vietnam/1203/2004 (H5N1)(13, 28) . In our hands, H5TX N224K Q226L bound human trachea sections, in contrast to H5TX Q226L. Unlike H5TX and H5TX Q226L, we observed binding of H5TX N224K Q226L to asymmetric α2,6-linked SIA on the glycan array, suggesting that the number and position of sialic acids could affect the receptor-binding properties of these H5TX mutants. This observation is consistent with previous findings that H3 binding to the human trachea correlated with interactions with asymmetrical N-glycans, specifically those featuring a single branch bearing a human-type sialic acid (25).

The HA protein derived from the 2.3.4.4e clade H5AK Q226L fully switches specificities from avian-type receptors to human-type receptors, not only on synthetic compounds but also from chicken to human trachea. To highlight the importance of evaluating each strain individually, we introduced H5AK key residues into the H5TX background. While position 227 has been linked to changes in H5 immunogenicity (29), changes in the 133 position have been associated to changes in receptor binding avidity and affinity across HA subtypes, including H1, H3, H5, and H7e (30-34). Single mutations did not affect either chicken or human tracheal binding, whereas combining them with Q226L resulted in a complete loss of tissue binding. Recovery of biantennary α2,6-linked SIA binding in both H5TX Δ133L Q226L and the triple mutant indicates that the Q226L mutation is insufficient to fully switch H5TX’s binding to human tracheal tissue sections, and that additional mutations are needed.

In this study, we also gained insights into how the presence of a multibasic cleavage site (MBCS) affects the receptor-binding properties of 2.3.4.4b hemagglutinins. We show that inserting an MBCS across a variety of strains does not generally affect receptor-binding properties. However, we observed that H5FR Q226L lost its binding to α2,3-linked SIA upon insertion of the MBCS. This finding suggests that the presence of an MBCS sequence can interfere with this particular HA’s receptor-binding specificity. Despite extensive studies of MBCS with respect to pathogenicity and virulence, it remains unknown whether these motifs play a role in receptor attachment. This study again shows the importance of checking each antigenic background individually, not only for point mutations but also for the effect of an MBCS on receptor-binding avidity and specificity. One suggestion for future research would be to analyze this H5FR Q226L (+) MBCS protein at the structural level to assess whether the insertion of MBCS affects folding and, if so, which bonds with the avian receptor analog are affected.

Differences in sensitivity among ELISA and glycan arrays highlight limitations in interpreting protein–glycan interactions (35). ELISA-based methods often offer signal amplification and can detect relatively low-affinity interactions, but they rely on immobilized glycans presented in a simplified, often non-native form. Glycan arrays offer higher throughput and greater structural diversity, yet the density, spatial arrangement, and overall repertoire of glycans remain highly restricted compared to those found in vivo, potentially leading to underestimation of binding breadth or affinity. These assays evaluate only a small subset of glycans and do not capture the full structural complexity, heterogeneity, and presentation of glycans on tissues such as the trachea. This simplification demonstrates the importance of complementing in vitro tests with tissue-based experiments.

Overall, we conclude that more than a single mutation is required for the hemagglutinin of the currently circulating 2.3.4.4b viruses to bind human respiratory tissue. This aligns with previous work showing that multiple amino acid changes are required to switch from avian-to human-type receptors in H5 strains (3). One key takeaway from this study is the significance of glycan architecture. H5TX mutants, similarly to other HAs distinguish not only between SIA linkage but are also selective according to glycan branch length and sialylation (25). More research is needed on the presence of asymmetrical glycans in the human respiratory tract and their role in influenza A virus hemagglutinin attachment to cells. Moreover, we emphasize the importance of not underestimating the MBCS’s influence on hemagglutinin’s receptor-binding properties.

## Materials and methods

### Expression and purification of hemagglutinin (change wording)

The pCD5 vector was engineered to allow in-frame insertion of HA cDNAs with sequences encoding a secretion signal, an N-terminal mOrange2 protein, a GCN4 trimerization domain (RMKQIEDKIEEIESKQKKIENEIARIKK), and a Twin-Strep tag (WSHPQFEKGGGSGGGSWSHPQFEK; IBA, Germany) (36-38). The resulting constructs were transfected into HEK293S GnTI(−) cells and purified as described previously (36). Briefly, DNA and polyethyleneimine I (PEI) were combined at a 1:8 ratio (µg DNA: µg PEI) and added to the cells. After 6 hours, the transfection mixture was replaced with 293 SFM II suspension medium (Invitrogen, 11686029) supplemented with glucose (2.0 g/L), sodium bicarbonate (3.6 g/L), primatone (3.0 g/L; Kerry), 1% glutaMAX (Gibco), 1.5% DMSO, and 2 mM valproic acid. Four days post-transfection, supernatants were collected, and proteins were purified using Strep-Tactin Sepharose beads (IBA Life Sciences) as previously described (39).

### Hemagglutination assay

Hemagglutination assay was performed as described previously (36). HA proteins were precomplexed with α-strep-tag human antibody (GenScript) and goat-α-mouse human antibody (Novus Biologicals) for 30 minutes at a molar ratio of 4:2:1, respectively. For nanoparticle assemblies, HA proteins were pooled with CompB in PBS overnight at 4 °C. Human and chicken erythrocytes were washed twice with PBS and diluted to 1% solution. Serial dilutions of protein mixes, precomplexed and NPs, were incubated for at least 1 hour at room temperature with red blood cells before visualization.

### Glycan array

The previously described glycan microarray was used (40). Hemagglutinins (HAs) were pre-complexed with mouse anti-Strep-tag-HRP and goat anti-mouse-Alexa555 antibodies at a molar ratio of 4:2:1 in 50 μL phosphate-buffered saline (PBS) containing 0.1% Tween-20. The mixture was incubated on ice for 20 minutes, then incubated on the array surface in a humidified chamber for 90 minutes. Slides were then washed sequentially with PBS-T (0.1% Tween-20), PBS, and deionized water. Arrays were dried by centrifugation and scanned immediately as previously described. For data processing, six replicates were analyzed by excluding the highest and lowest values, and calculating the mean and standard deviation from the remaining four replicates.

### Protein histochemistry (Tissue staining)

Histochemical analysis was performed as previously described (41). Paraffin-embedded tissue sections from human and chicken trachea were re-hydrated with 100%, 95%,70% ethanol and distilled water. Antigens were retrieved with warm citric acid at pH 6.0, and endogenous peroxidase was blocked with 1% H_2_0_2_ in methanol for 20 minutes at RT. Tissues were washed with PBS-Tween and incubated in 3% BSA overnight at 4 °C. HA proteins were precomplexed with α-strep-tag human antibody (GenScript) and goat-α-mouse human antibody (Novus Biologicals) for 30 minutes at a molar ratio of 4:2:1, respectively. For nanoparticle assemblies, HA proteins were pooled with CompB in PBS overnight at 4 °C. Protein mixes were added and incubated for at least 90 minutes at room temperature. Slides were then washed with PBS and incubated with DAPI for 10 minutes in the dark, then mounted with FluorSave (). Images were taken using a Leica DMi8 confocal microscope equipped with a 10× HC PL Apo CS2 objective (NA 0.40) as described before (42). Image analysis was performed using LAS Application Suite X and ImageJ.

### Glycan ELISA

Glycan ELISA assay was performed as previously described (25). Nunc Maxisorp 96-well plates (Invitrogen) were coated overnight at 4°C with 50 μL of 5 μg/mL streptavidin (Westburg) in PBS. Plates were then blocked with 300 μL of 1% BSA in PBS-T for 3 hours at room temperature, followed by incubation with 50 μL of 50 nM biotinylated glycans overnight at 4°C. After a second blocking step with 300 μL of 1% BSA in PBS-T for 3 hours at room temperature, NP mixtures were added, serially diluted 1:3, and incubated for 90 minutes at room temperature.

Detection was carried out using an OPD substrate solution, and the reaction was stopped after 5 minutes with 2.5 M H_2_SO_4_. Absorbance was measured at 485 nm using a Polar Star Omega plate reader (BMG Labtech), and raw data were analyzed in GraphPad Prism.

## Acknowledgements

M.R.C is supported by an NWO-M2 (OCENW.M20.106). This research was made possible by funding from ICRAD, an ERA-NET co-funded under the European Union’s Horizon 2020 research and innovation programme (https://ec.europa.eu/programmes/horizon2020/en), under Grant Agreement no. 862605 (Flu-Switch).

## References

1. Lewis NS, Banyard AC, Whittard E, Karibayev T, Al Kafagi T, Chvala I, Byrne A, Meruyert Akberovna S, King J, Harder T, Grund C, Essen S, Reid SM, Brouwer A, Zinyakov NG, Tegzhanov A, Irza V, Pohlmann A, Beer M, Fouchier RAM, Akhmetzhan Akievich S, Brown IH. 2021. Emergence and spread of novel H5N8, H5N5 and H5N1 clade 2.3.4.4 highly pathogenic avian influenza in 2020. Emerg Microbes Infect 10:148–151.

2. Plaza PI, Gamarra-Toledo V, Eugui JR, Lambertucci SA. 2024. Recent Changes in Patterns of Mammal Infection with Highly Pathogenic Avian Influenza A(H5N1) Virus Worldwide. Emerg Infect Dis 30:444–452.

3. Paulson JC, de Vries RP. 2013. H5N1 receptor specificity as a factor in pandemic risk. Virus Res 178:99–113.

4. Santos JJS, Wang S, McBride R, Adams L, Harvey R, Zhao Y, Wrobel AG, Gamblin S, Skehel J, Lewis NS, Paulson JC, Hensley SE. 2025. Bovine H5N1 binds poorly to human-type sialic acid receptors. Nature 640:E18–E20.

5. Zhu B, Fung K, Feng HH, Beatty JA, Hill F, Tse ACN, Brackman CJ, Sit THC, Poujade A, Gaide N, Ducatez M, Foucras G, Peiris M, Ti SC, Nicholls JM, Yen HL. 2025. The hemagglutinin proteins of clades 1 and 2.3.4.4b H5N1 highly pathogenic avian influenza viruses exhibit comparable attachment patterns to avian and mammalian tissues. J Virol 99:e0097625.

6. Chopra P, Page CK, Shepard JD, Ray SD, Kandeil A, Jeevan T, Bowman AS, Ellebedy AH, Webby RJ, de Vries RP, Tompkins SM, Boons GJ. 2024. Receptor Binding Specificity of a Bovine A(H5N1) Influenza Virus. bioRxiv doi:10.1101/2024.07.30.605893.

7. Good MR, Fernandez-Quintero ML, Ji W, Rodriguez AJ, Han J, Ward AB, Guthmiller JJ. 2024. A single mutation in dairy cow-associated H5N1 viruses increases receptor binding breadth. Nat Commun 15:10768.

8. Kamel M, Aleya S, Almagharbeh WT, Aleya L, Abdel-Daim MM. 2025. The emergence of highly pathogenic avian influenza H5N1 in dairy cattle: implications for public health, animal health, and pandemic preparedness. Eur J Clin Microbiol Infect Dis 44:1817–1833.

9. Mostafa A, Naguib MM, Nogales A, Barre RS, Stewart JP, Garcia-Sastre A, Martinez-Sobrido L. 2024. Avian influenza A (H5N1) virus in dairy cattle: origin, evolution, and cross-species transmission. mBio 15:e0254224.

10. Rios Carrasco M, Lin TH, Zhu X, Gabarroca Garcia A, Uslu E, Liang R, Spruit CM, Richard M, Boons GJ, Wilson IA, de Vries RP. 2025. The Q226L mutation can convert a highly pathogenic H5 2.3.4.4e virus to bind human-type receptors. Proc Natl Acad Sci U S A 122:e2419800122.

11. Lin TH, Zhu X, Wang S, Zhang D, McBride R, Yu W, Babarinde S, Paulson JC, Wilson IA. 2024. A single mutation in bovine influenza H5N1 hemagglutinin switches specificity to human receptors. Science 386:1128–1134.

12. Eggink D, Spronken M, van der Woude R, Buzink J, Broszeit F, McBride R, Pawestri HA, Setiawaty V, Paulson JC, Boons GJ, Fouchier RAM, Russell CA, de Jong MD, de Vries RP. 2020. Phenotypic Effects of Substitutions within the Receptor Binding Site of Highly Pathogenic Avian Influenza H5N1 Virus Observed during Human Infection. J Virol 94.

13. Imai M, Watanabe T, Hatta M, Das SC, Ozawa M, Shinya K, Zhong G, Hanson A, Katsura H, Watanabe S, Li C, Kawakami E, Yamada S, Kiso M, Suzuki Y, Maher EA, Neumann G, Kawaoka Y. 2012. Experimental adaptation of an influenza H5 HA confers respiratory droplet transmission to a reassortant H5 HA/H1N1 virus in ferrets. Nature 486:420–8.

14. Peng W, Bouwman KM, McBride R, Grant OC, Woods RJ, Verheije MH, Paulson JC, de Vries RP. 2018. Enhanced Human-Type Receptor Binding by Ferret-Transmissible H5N1 with a K193T Mutation. J Virol 92.

15. West J, Roder J, Matrosovich T, Beicht J, Baumann J, Mounogou Kouassi N, Doedt J, Bovin N, Zamperin G, Gastaldelli M, Salviato A, Bonfante F, Kosakovsky Pond S, Herfst S, Fouchier R, Wilhelm J, Klenk HD, Matrosovich M. 2021. Characterization of changes in the hemagglutinin that accompanied the emergence of H3N2/1968 pandemic influenza viruses. PLoS Pathog 17:e1009566.

16. Schrauwen EJA, Bestebroer TM, Munster VJ, de Wit E, Herfst S, Rimmelzwaan GF, Osterhaus A, Fouchier RAM. 2011. Insertion of a multibasic cleavage site in the haemagglutinin of human influenza H3N2 virus does not increase pathogenicity in ferrets. J Gen Virol 92:1410–1415.

17. Suguitan AL, Jr., Matsuoka Y, Lau YF, Santos CP, Vogel L, Cheng LI, Orandle M, Subbarao K. 2012. The multibasic cleavage site of the hemagglutinin of highly pathogenic A/Vietnam/1203/2004 (H5N1) avian influenza virus acts as a virulence factor in a host-specific manner in mammals. J Virol 86:2706–14.

18. Abdelwhab EM, Veits J, Ulrich R, Kasbohm E, Teifke JP, Mettenleiter TC. 2016. Composition of the Hemagglutinin Polybasic Proteolytic Cleavage Motif Mediates Variable Virulence of H7N7 Avian Influenza Viruses. Sci Rep 6:39505.

19. Abdelwhab el SM, Veits J, Tauscher K, Ziller M, Teifke JP, Stech J, Mettenleiter TC. 2016. A Unique Multibasic Proteolytic Cleavage Site and Three Mutations in the HA2 Domain Confer High Virulence of H7N1 Avian Influenza Virus in Chickens. J Virol 90:400–11.

20. Broszeit F, van Beek RJ, Unione L, Bestebroer TM, Chapla D, Yang JY, Moremen KW, Herfst S, Fouchier RAM, de Vries RP, Boons GJ. 2021. Glycan remodeled erythrocytes facilitate antigenic characterization of recent A/H3N2 influenza viruses. Nat Commun 12:5449.

21. Ma S, Liu L, Eggink D, Herfst S, Fouchier RAM, de Vries RP, Boons GJ. 2024. Asymmetrical Biantennary Glycans Prepared by a Stop-and-Go Strategy Reveal Receptor Binding Evolution of Human Influenza A Viruses. JACS Au 4:607–618.

22. Fabrizio TP, Kandeil A, Harrington WN, Jones JC, Jeevan T, Andreev K, Seiler P, Fogo J, Davis ML, Crumpton JC, Franks J, DeBeauchamp J, Vogel P, Daniels CS, Poulson RL, Bowman AS, Govorkova EA, Webby RJ. 2025. Genotype B3.13 influenza A(H5N1) viruses isolated from dairy cattle demonstrate high virulence in laboratory models, but retain avian virus-like properties. Nat Commun 16:6771.

23. Yang J, Qureshi M, Kolli R, Peacock TP, Sadeyen JR, Carter T, Richardson S, Daines R, Barclay WS, Brown IH, Iqbal M. 2025. The haemagglutinin gene of bovine-origin H5N1 influenza viruses currently retains receptor-binding and pH-fusion characteristics of avian host phenotype. Emerg Microbes Infect 14:2451052.

24. Weber J, Ponse NLD, Zhu X, Rios Carrasco M, Han AX, Funk M, Lin TH, Gabarroca Garcia A, Spruit CM, Zhang D, Yu W, Wilson IA, Richard M, Boons GJ, de Vries RP. 2026. The receptor binding properties of H5Ny influenza A viruses have evolved to bind to avian-type mucin-like O-glycans. PLoS Pathog 22:e1013812.

25. Liang R, Peccati F, Ponse NLD, Uslu E, de Rooij AJH, Han AX, Boons GJ, Unione L, de Vries RP. 2025. Epistasis in the receptor-binding domain of contemporary H3N2 viruses that reverted to bind sialylated di-LacNAc repeats. Cell Rep 44:116007.

26. Bauer L, Leijten L, Iervolino M, Chopra V, van Dijk L, Power M, Rijnink W, Pronk M, Spronken M, Funk M, de Vries RD, Richard M, Kuiken T, van Riel D. 2026. Attachment and replication of clade 2.3.4.4b influenza A (H5N1) viruses in human respiratory epithelium: an in-vitro study. Lancet Microbe 7:101230.

27. Yu H, Gonzalez-Gil A, Wei Y, Fernandes SM, Porell RN, Vajn K, Paulson JC, Nycholat CM, Schnaar RL. 2017. Siglec-8 and Siglec-9 binding specificities and endogenous airway ligand distributions and properties. Glycobiology 27:657–668.

28. Dadonaite B, Ahn JJ, Ort JT, Yu J, Furey C, Dosey A, Hannon WW, Vincent Baker AL, Webby RJ, King NP, Liu Y, Hensley SE, Peacock TP, Moncla LH, Bloom JD. 2024. Deep mutational scanning of H5 hemagglutinin to inform influenza virus surveillance. PLoS Biol 22:e3002916.

29. Hu Z, Shi L, Zhao J, Gu H, Hu J, Wang X, Liu X, Hu S, Gu M, Cao Y, Liu X. 2020. Role of the Hemagglutinin Residue 227 in Immunogenicity of H5 and H7 Subtype Avian Influenza Vaccines in Chickens. Avian Dis 64:445–450.

30. Koerner I, Matrosovich MN, Haller O, Staeheli P, Kochs G. 2012. Altered receptor specificity and fusion activity of the haemagglutinin contribute to high virulence of a mouse-adapted influenza A virus. J Gen Virol 93:970–979.

31. Simmons HC, Watanabe A, Oguin Iii TH, Van Itallie ES, Wiehe KJ, Sempowski GD, Kuraoka M, Kelsoe G, McCarthy KR. 2023. A new class of antibodies that overcomes a steric barrier to cross-group neutralization of influenza viruses. PLoS Biol 21:e3002415.

32. Tharakaraman K, Raman R, Viswanathan K, Stebbins NW, Jayaraman A, Krishnan A, Sasisekharan V, Sasisekharan R. 2013. Structural determinants for naturally evolving H5N1 hemagglutinin to switch its receptor specificity. Cell 153:1475–85.

33. Wang Y, Li Q, Peng P, Zhang Q, Huang Y, Hu J, Hu Z, Liu X. 2024. Dual N-linked glycosylation at residues 133 and 158 in the hemagglutinin are essential for the efficacy of H7N9 avian influenza virus like particle vaccine in chickens and mice. Vet Microbiol 294:110108.

34. Zhu X, Viswanathan K, Raman R, Yu W, Sasisekharan R, Wilson IA. 2015. Structural Basis for a Switch in Receptor Binding Specificity of Two H5N1 Hemagglutinin Mutants. Cell Rep 13:1683–91.

35. Pochechueva T, Jacob F, Goldstein DR, Huflejt ME, Chinarev A, Caduff R, Fink D, Hacker N, Bovin NV, Heinzelmann-Schwarz V. 2011. Comparison of printed glycan array, suspension array and ELISA in the detection of human anti-glycan antibodies. Glycoconj J 28:507–17.

36. de Vries RP, de Vries E, Bosch BJ, de Groot RJ, Rottier PJ, de Haan CA. 2010. The influenza A virus hemagglutinin glycosylation state affects receptor-binding specificity. Virology 403:17–25.

37. Harbury PB, Zhang T, Kim PS, Alber T. 1993. A switch between two-, three-, and four-stranded coiled coils in GCN4 leucine zipper mutants. Science 262:1401–7.

38. Sliepen K, van Montfort T, Ozorowski G, Pritchard LK, Crispin M, Ward AB, Sanders RW. 2015. Engineering and Characterization of a Fluorescent Native-Like HIV-1 Envelope Glycoprotein Trimer. Biomolecules 5:2919–34.

39. Krammer F, Margine I, Tan GS, Pica N, Krause JC, Palese P. 2012. A carboxy-terminal trimerization domain stabilizes conformational epitopes on the stalk domain of soluble recombinant hemagglutinin substrates. PLoS One 7:e43603.

40. Broszeit F, Tzarum N, Zhu X, Nemanichvili N, Eggink D, Leenders T, Li Z, Liu L, Wolfert MA, Papanikolaou A, Martinez-Romero C, Gagarinov IA, Yu W, Garcia-Sastre A, Wennekes T, Okamatsu M, Verheije MH, Wilson IA, Boons GJ, de Vries RP. 2019. N-Glycolylneuraminic Acid as a Receptor for Influenza A Viruses. Cell Rep 27:3284–3294 e6.

41. Nemanichvili N, Berends AJ, Wubbolts RW, Grone A, Rijks JM, de Vries RP, Verheije MH. 2021. Tissue Microarrays to Visualize Influenza D Attachment to Host Receptors in the Respiratory Tract of Farm Animals. Viruses 13.

42. Tomris I, Unione L, Nguyen L, Zaree P, Bouwman KM, Liu L, Li Z, Fok JA, Rios Carrasco M, van der Woude R, Kimpel ALM, Linthorst MW, Kilavuzoglu SE, Verpalen E, Caniels TG, Sanders RW, Heesters BA, Pieters RJ, Jimenez-Barbero J, Klassen JS, Boons GJ, de Vries RP. 2023. SARS-CoV-2 Spike N-Terminal Domain Engages 9-O-Acetylated alpha2-8-Linked Sialic Acids. ACS Chem Biol 18:1180–1191.

